# Nutrient cycling is an important mechanism for homeostasis in plant cells

**DOI:** 10.1101/2021.04.01.438083

**Authors:** Ingo Dreyer

**Author notes:** **Footnotes Author contributions:** ID conceived the project, designed the experiments, analyzed the data, prepared the figures and wrote the article. **Funding:** This research did not receive any specific grant from funding agencies in the public, commercial, or not-for-profit sectors.

## Abstract

Homeostasis in living cells refers to the steady state of internal, physical, and chemical conditions. It is maintained by self-regulation of the dynamic cellular system. In order to gain insight into homeostatic mechanisms that keep cytosolic nutrient concentrations in plant cells within a homeostatic range, I performed computational cell biology experiments. Systems of membrane transporters were modelled mathematically followed by the simulation of their dynamics. The detailed analyses of ‘*what-if*’ scenarios demonstrate that a single transporter type for a nutrient, irrespective whether it is a channel or a co-transporter, is not sufficient to set a desired cytosolic concentration. A cell cannot flexibly react on different external conditions. At least two different transporter types for the same nutrient are required, which are energized differently. The gain of flexibility in adjusting the nutrient concentration was accompanied by the establishment of energy-consuming nutrient cycles at the membrane suggesting that these sometimes called ‘futile’ cycles are not as futile as they appear. This understanding may help in future to design new strategies for increasing nutrient use efficiency of crop plants taking into account the complex interplay of transporter networks at the cellular level.

**One sentence summary:** First principles of membrane transport explain why the maintenance of a constant cytosolic nutrient concentration is often accompanied by the ‘futile’ cycling of the nutrient across the membrane.

## Introduction

For plants as for animals, control of (ion) homeostasis is crucial to establish defined conditions in cells and cellular compartments that finally allow various biochemical processes to take place in a controlled manner. Homeostatic processes are therefore fundamental for plant life development and resilience to even hostile environments. A critical role in (ion) homeostasis is played by membrane transport proteins. As estimated for Arabidopsis, around 18% of the predicted proteins of a plant genome may contain two or more transmembrane domains and about more than half of those may perform transporter functions (Ward, 2001; Schwacke et al., 2003; Tang et al., 2020). Although our knowledge on the function, the tissue-specific expression and subcellular localization of plant membrane transporters increases daily, detailed mechanistic insights into the functional interplay between these transporter proteins are still largely missing. This gap in knowledge might be a severe obstacle in developing new strategies and approaches involving transporter proteins to improve nutrient use efficiency and tolerance to environmental stresses as well as to boost general crop fitness and productivity.

The analysis of membrane transporter networks is not a trivial task and almost impossible to accomplish with exclusively wet-laboratory approaches. Nevertheless, transporter types can be mathematically described, allowing deep insights into the dynamics of transporter networks by computational analyses (Dreyer and Michard, 2020). Computational modelling helped already to better understand guard cell movement (Hills et al., 2012; Blatt et al., 2014), the role of K^+^ gradients in energizing phloem (re-)loading processes (Gajdanowicz et al., 2011; Dreyer et al., 2017), the nutrient exchange in mycorrhizal symbioses (Schott et al., 2016; Dreyer et al., 2019; Nizam et al., 2019), the mechanism and potential consequences of vacuolar excitability (Jaslan et al., 2019; Dindas et al., 2021) and touch-sensitive signaling in trigger hairs of the Venus flytrap (Iosip et al., 2020).

The present study was inspired by the efforts to optimize K^+^ use efficiency in crop plants (Zörb et al., 2014). A large variety of K^+^ transporters has been identified that are employed by plants for the task of K^+^ uptake and redistribution. Additionally, our knowledge on the signaling cascades that regulate the activity of these transporters is growing daily (for reviews, see e.g. Dreyer and Uozumi, 2011; Sharma et al., 2013; Ahmad and Maathuis, 2014; Anschütz et al., 2014; Chérel et al., 2014; Nieves-Cordones et al., 2014; Shabala and Pottosin, 2014; Véry et al., 2014; Luan et al., 2017; Sze and Chanroj, 2018; Chérel and Gaillard, 2019; Ragel et al., 2019).

Nevertheless, it is still not clear which constraints set limits to a cell to control the cytosolic K^+^ concentration by adjusting the transporter activities in response to the external conditions. To address this knowledge gap, in this study computational cell biology experiments have been designed that provide comprehensible insight into a cell’s ability to control cytosolic [K^+^] under given external conditions. The topic is approached in thought experiments (*‘what-if’* scenarios) to illustrate the straightforward conclusions from thermodynamic basics of membrane transport and mathematical considerations. This approach allowed to understand in general fundamental principles of homeostasis that permit a living compartment (a cell or an organelle) to set defined internal conditions. Although the motivating starting point was K^+^ nutrition, the results were be generalized to other nutrients. The conclusions presented here provide fundamental insights into the interplay between different transporters and expose essential features needed to control homeostasis. The new insights might provide important theoretical knowledge for future attempts to optimize nutrient use efficiency in crop plants.

## Results

The goal of this study was to answer the question, in as much can a plant cell adjust the steady state of internal concentrations and the membrane voltage by changing membrane transporter activities. This is not a trivial problem and can easily reach a complexity that goes beyond our imagination. To gain insights nevertheless, a step-wise synthetic computational cell biology approach was chosen. Starting from the analysis of the simplest systems, the complexity was successively increased in order to learn how this affects the flexibility of the systems. In a first set of thought experiments, a cell was regarded that is surrounded only by the plasma membrane. *I*.*e*., at that stage the effects introduced by organelles like the vacuole were eliminated. This type of complexity was added in a subsequent step. The conceptional approach is presented first for the case of the most abundant cation in plants, K^+^, followed by counter charge providing anions and finally for sugar as an example for an uncharged nutrient and for metabolic processes.

### The control of [K^+^]_cyt_ implies the control of E_K_

An important note at the beginning: Membrane transport processes are ruled by the transmembrane (electro)chemical gradient of the considered nutrient. This gradient does not depend solely on the cytosolic concentration (e.g. [K^+^]_cyt_), but on the logarithm of the ratio of the concentrations on both sides of the membrane 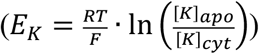. In the case of potassium, a cell can primarily control *E*_*K*_ by adjusting the activities of the membrane transporters. In turn, through the control of *E*_*K*_, [K^+^]_cyt_ can be controlled and maintained constant in the case of homeostasis. In other words, in order to respond to a wider range of external conditions when holding [K^+^]_cyt_ constant, a cell must be able to adjust *E*_*K*_ over a wider range.

### Case 1: only K^+^ channels

A first *‘what-if’* scenario considered the case, in which the proton H^+^-ATPase acted together with K^+^ channels. In such a system, the H^+^-ATPase pumped H^+^ out of the cell, while the open potassium channel allowed the influx of K^+^ that electrically compensated the H^+^ efflux (**Fig. 1 A**,**B**). The accumulation of K^+^ in the cell led to a negative shift of *E*_*K*_, which also negatively shifted the voltage, at which the H^+^ efflux compensated the K^+^ influx. A stable steady state could only be reached in this system if *I*_*P*_(*V*) = 0 and *E*_*K*_ = *V*. This means that the membrane voltage adjusted to the steady state voltage of the H^+^-ATPase and that also *E*_*K*_ was fixed to this value.

**Figure 1.**
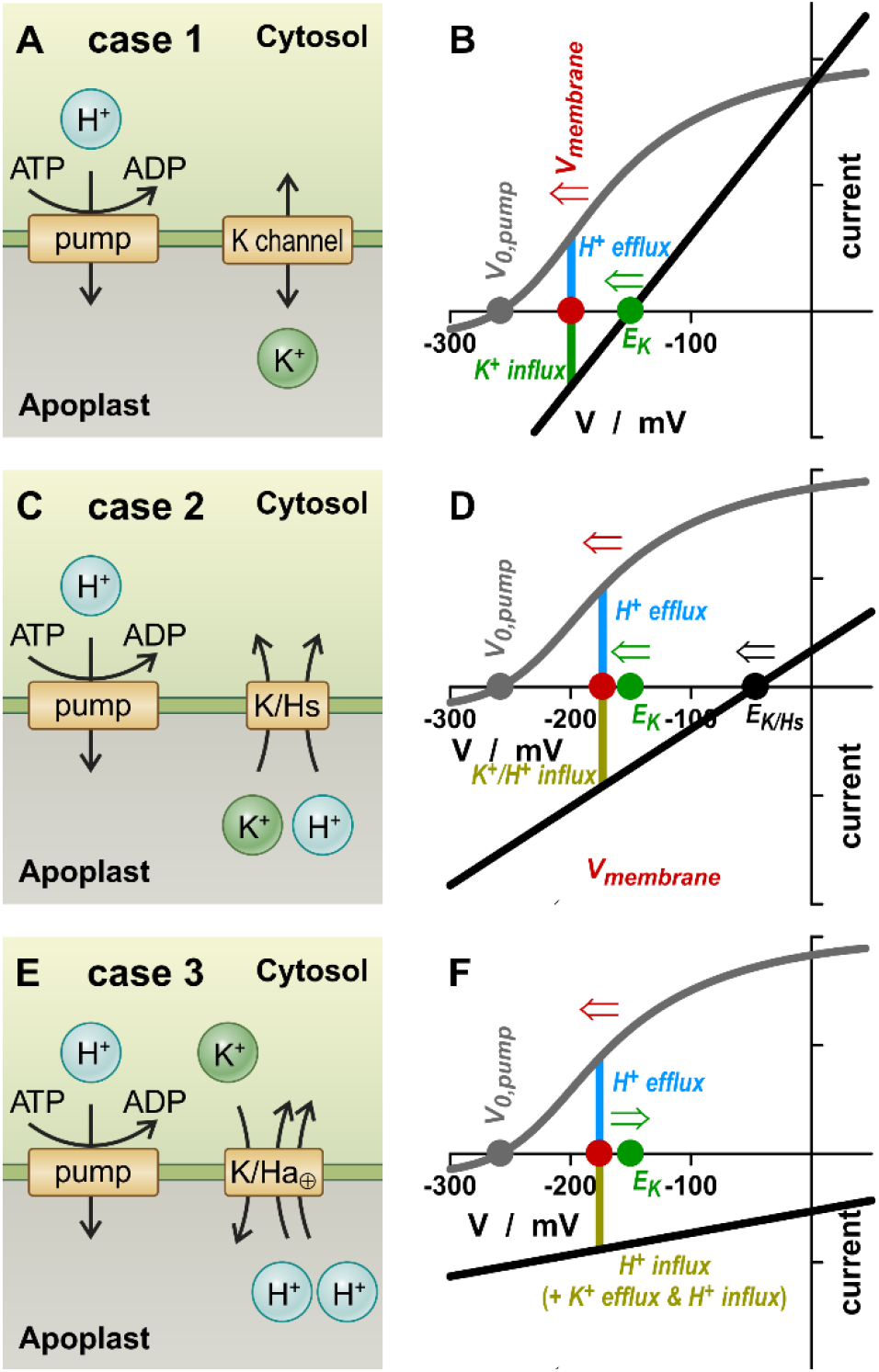
*E*_*K*_ and *V* adjusted by single K^+^ transporters and the H^+^ pump. (A,B) Case 1, only K^+^ channels. The H^+^ efflux is electrically compensated by a K^+^ influx. The only stable steady state of the system is *E*_*K*_ = *V*_*0,pump*_. (C,D) Case 2, only K^+^/H^+^ symporters. The H^+^ efflux is electrically compensated by the K^+^/H^+^ influx. The only stable steady state of the system is *E*_*K*_ = 2·*V*_*0,pump*_ – *E*_*H*_. (E,F) Case 3, only K^+^/H^+^ antiporters. In case of electrogenic K^+^/2H^+^ antiporters, the pump-mediated H^+^ efflux is electrically compensated by the H^+^-mediate charge surplus of the antiport. In general the steady state of this system is *E*_*K*_ = *n*·*E*_*H*_ – (*n*-1)·*V*_*0,pump*_.

There was neither a net K^+^ flux via the channel nor a net H^+^ flux via the pump under this condition. To get an idea, which K^+^ gradients could be established with such a system, a normal pH gradient (pH_cyt_ = 7.0, pH_apo_ = 6.0) and a steady state voltage of *V*_*0,pump*_ = -200 mV were considered. For an external K^+^ concentration of [K^+^]_apo_ = 100 μM the internal concentration would have equilibrated at [K^+^]_cyt_ = 300 mM, and for [K^+^]_apo_ = 1 mM the steady state had been at [K^+^]_cyt_ = 3 M. These cation concentrations would need to be charge-balanced by anions, for instance malate^2-^ originating from metabolic processes that also provide the protons pumped out of the cell. The consequences of anion transport and metabolic processes are considered further below. The steady state of case 1 was determined exclusively by the voltage, at which the pump current was zero [parameter *V*_*0,pump*_ in eqn (1)]. This parameter depends on the energetic status of the cell and the transmembrane pH-gradient. If, besides pump and K^+^ channel, there were other transporters in the membrane that were not involved in K^+^ transport, they would consume part of the energy provided by the H^+^-ATPase. As it will be shown later in this article, such a situation can be simulated in the considered steady state model simply by adjusting the parameter *V*_*0,pump*_.

With *V*_*0,pump*_ = -125 mV the results were qualitatively identical as with *V*_*0,pump*_ = -200 mV. However, *E*_*K*_ would now be -125 mV in the steady state and the K^+^ gradients [K^+^]_apo_ = 100 μM / [K^+^]_cyt_ = 14.8 mM and [K^+^]_apo_ = 1 mM / [K^+^]_cyt_ = 148 mM, respectively. Thus, the cell can control *E*_*K*_ and [K^+^]_cyt_ indirectly via the energy not dissipated elsewhere, therefore available for K^+^ transport. However, a direct control of *V* or *E*_*K*_ by varying the K^+^ channel activity was not possible.

### Case 2: only K^+^/H^+^ symporters

A second *‘what-if’* scenario considered the case, in which the proton H^+^-ATPase acted together with K^+^/H^+^ symporters. Here, the pumped H^+^ were electrically compensated by a combined K^+^/H^+^ influx (**Fig. 1C**,**D**). The net K^+^ influx shifted *E*_*K*_ to more negative voltages and thus also the zero-current voltage of the symporter, *E*_*K/Hs*_. This shifted the membrane voltage, at which the positive and negative electrical fluxes compensated each other, to more negative values. Similar to case 1, the system had only one possible steady state determined by the equilibrium voltage of the H^+^-ATPase: *I*_*P*_(*V*) = 0, which implied *V* = *V*_*0,pump*_, and *E*_*K*_ = 2·*V*_*0,pump*_–*E*_*H*_. Under this condition, the K^+^/H^+^ flux via the symporter was zero. With pH_cyt_ = 7.0, pH_apo_ = 6.0, and *V*_*0,pump*_ = -200 mV, the internal concentration would have equilibrated in this system at [K^+^]_cyt_ = 9×10^4^ mM for [K^+^]_apo_ = 1 μM, and at [K^+^]_cyt_ = 9×10^7^ mM for [K^+^]_apo_ = 1 mM. Of course, these values are purely theoretical and are intended to illustrate how far the modeled scenario is from a physiological situation. The exorbitant gradients declined to still huge [K^+^]_apo_ = 1 μM / [K^+^]_cyt_ = 220 mM and [K^+^]_apo_ = 1 mM / [K^+^]_cyt_ = 2.2×10^5^ mM, respectively, when setting *V*_*0,pump*_ = -125 mV in order to simulate energy consumption by other transport processes (see case 1). Also here, a control of *V* or *E*_*K*_ by varying the K^+^/H^+^ symporter activity was not possible. Thus, a system of pump and K^+^/H^+^ symporter is even more susceptible to K^+^ over-accumulation than a “pump & channel” system (case 1).

### Case 3: only K^+^/H^+^ antiporters

A third *‘what-if’* scenario considered the case, in which the proton H^+^-ATPase acted together with K^+^/H^+^ antiporters, which exchanged one K^+^ for *n*H^+^. An electroneutral K^+^/H^+^ antiport was represented by *n* = 1 and an electrogenic antiport by *n* > 1. The system with an electroneutral K^+^/H^+^ antiporter released K^+^ until the K^+^ gradient equaled the H^+^ gradient (*E*_*K*_ = *E*_*H*_). This release was energized by the proton gradient. The steady state membrane voltage was *V*_*0,pump*_, at which no net ion flux occurred. In case of an electrogenic K^+^/H^+^ antiporter, the released K^+^ ion was electrically compensated by one inflowing H^+^, while the other protons transported by the antiporter were electrically compensated by protons pumped out of the cell by the H^+^-ATPase. The net proton fluxes were compensated by biochemical/metabolic processes (Sanders and Slayman, 1982; Wegner and Shabala, 2020; Wegner et al., 2021), whereas the release of K^+^ shifted *E*_*K*_ positively along the voltage axis (**Fig. 1E**,**F**). A steady state was only achieved if *I*_*P*_(*V*) = 0, which implied *V* = *V*_*0,pump*_ and *E*_*H/Ka*_ = *V*_*0,pump*_. Thus, in steady state was *E*_*K*_ = *n*·*E*_*H*_–(*n*-1)·*V*_*0,pump*_. With this value, the K^+^/H^+^ flux via the antiporter was zero at *V*_*0,pump*_. Considering the hypothetical case *n* = 2, the internal concentration would have equilibrated for pH_cyt_ = 7.0, pH_apo_ = 6.0, and *V*_*0,pump*_ = -200 mV at [K^+^]_cyt_ = 3.4 × 10^−9^ mM for [K^+^]_apo_ = 1 μM, and at [K^+^]_cyt_ = 3.4 × 10^−6^ mM for [K^+^]_apo_ = 1 mM. In an example of energy consumption by other transport processes (simulated by *V*_*0,pump*_ = -125 mV) the gradients would be [K^+^]_apo_ = 1 μM / [K^+^]_cyt_ = 6.7 × 10^−8^ mM and [K^+^]_apo_ = 1 mM / [K^+^]_cyt_ = 6.7 × 10^−5^ mM, respectively. As in the cases before, a control of *V* or *E*_*K*_ by varying the K^+^/H^+^ antiporter activity was not possible.

### A first interim conclusion

The considerations presented in cases 1-3 allowed already a very fundamental conclusion: An isolated, static cell could not flexibly adjust *E*_*K*_ with a single K^+^ transporter type. The steady state was always determined by the type of the transport process, the transmembrane pH gradient, and the H^+^-ATPase-dependent energy available for the K^+^ transport process (represented by *V*_*0,pump*_ in the model). The steady state varied between extremely low and extremely high unphysiological values indicating that these simple K^+^ transport modules are not (or only conditionally) suitable for a well-regulated K^+^ homeostasis at the plasma membrane. In the following it was examined how a mixture of different K^+^ transporters can increase the cellular flexibility.

### Case 4: electroneutral and electrogenic K^+^/H^+^ antiporters

In a fourth *‘what-if’* scenario the proton H^+^-ATPase was combined with both, electroneutral and electrogenic K^+^/H^+^ antiporters at the same time. The system established a steady state, in which the H^+^-ATPase pumped protons out of the cell, the electrogenic K^+^/H^+^ antiporter allowed the influx of two or more protons for the efflux of a K^+^ ion, while the electroneutral K^+^/H^+^ antiporter released a proton for the uptake of a K^+^ ion (**Fig. 2A**). Thus, apparently futile K^+^ and H^+^ cycles dissipated valuable energy provided by ATP-hydrolysis at the pump. However, the cell could now adjust *E*_*K*_ and the membrane voltage by fine-tuning the transporter activity (**Fig. 2B**,**C**).

**Figure 2.**
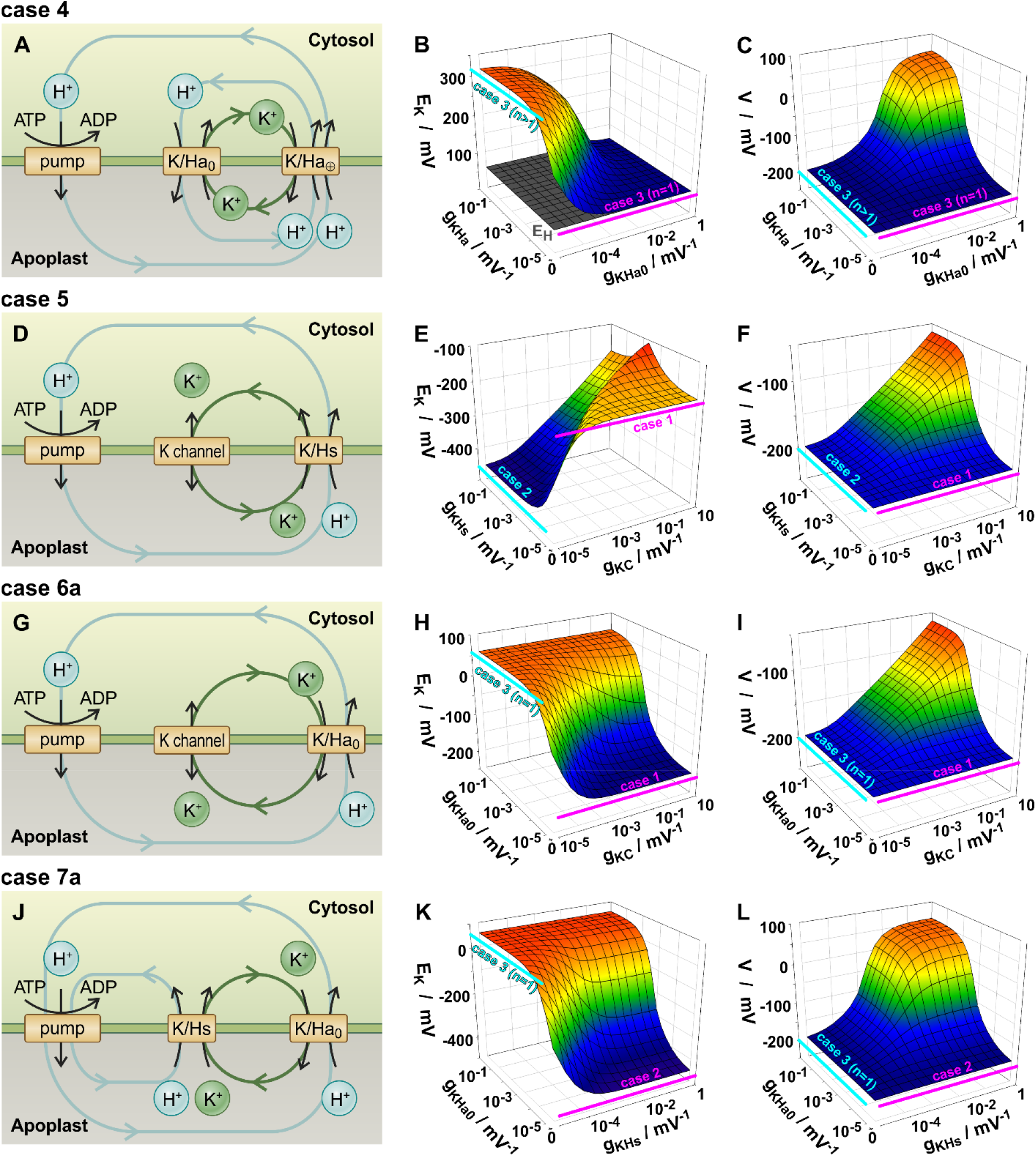
*E*_*K*_ and *V* adjusted by pairs of K^+^ transporters and the H^+^ pump. (A-C) Case 4. When electroneutral and electrogenic K^+^/H^+^ antiporters act together with the H^+^-pump, three cycles establish in steady state: (i) the electrogenic K^+^/H^+^ antiporter releases K^+^, which is reabsorbed by the electroneutral antiporter; (ii & iii) a proton is released by the electroneutral antiporter (ii), which is reabsorbed by the electrogenic antiporter together with the H^+^ released by the pump (iii). By modifying the transporter activities, the steady state of *E*_*K*_ and *V* can be freely adjusted in a certain range. For comparison, the respective steady states for the systems with only one K^+^ transporter-type: only electrogenic K^+^/H^+^ antiporters (*g*_*KHa0*_ = 0; Case 3, *n* > 1, cyan lines) and only electroneutral K^+^/H^+^ antiporters (*g*_*KHa*_ = 0; Case 3, *n* = 1, magenta lines). (D-F) Case 5a. When K^+^ channels and K^+^/H^+^ symporters act together with the H^+^-pump, two cycles establish in steady state: (i) the K^+^ channel releases K^+^, which is reabsorbed by the symporter together with the H^+^ released by the pump (ii). By modifying the transporter activities, the steady state of *E*_*K*_ and *V* can be freely adjusted in a certain range. For comparison, the respective steady states for the systems with only one K^+^ transporter-type: only K^+^/H^+^ symporters (*g*_*KC*_ = 0; Case 2, cyan lines) and just K^+^ channels (*g*_*KHs*_ = 0; Case 1, magenta lines). (G-I) Case 6a. When K^+^ channels and K^+^/H^+^ antiporters act together with the H^+^-pump, two cycles establish in steady state: (i) the K^+^ channel absorbs K^+^, which is released by the antiporter; (ii) the H^+^ released by the pump is absorbed by the antiporter. By modifying the transporter activities, the steady state of *E*_*K*_ and *V* can be freely adjusted in a certain range. For comparison, the respective steady states for the systems with only one K^+^ transporter-type: only K^+^/H^+^ antiporters (*g*_*KC*_ = 0; Case 3, *n* = 1, cyan lines) and only K^+^ channels (*g*_*KHa*0_ = 0; Case 1, magenta lines). (J-L) Case 7a. When K^+^/H^+^ symporters and K^+^/H^+^ antiporters act together with the H^+^-pump, three cycles establish in steady state: (i) the symporter absorbs K^+^, which is released by the antiporter; (ii) a proton released by the pump is absorbed by the symporter, (iii) while another released proton is absorbed by the antiporter. By modifying the transporter activities, the steady state *E*_*K*_ and *V* can be freely adjusted in a certain range. For comparison, the respective steady states for the systems with only one K^+^ transporter-type: only K^+^/H^+^ antiporters (*g*_*KHs*_ = 0; Case 3, *n* = 1, cyan lines) and only K^+^/H^+^ symporters (*g*_*KHa*0_ = 0; Case 2, magenta lines).

A comprehensive analysis of the steady state condition revealed that compared to the cases 1-3, which were characterized by exactly one steady state for *E*_*K*_ and *V* irrespective of the transporter activities (**Fig. 3A**), the case 4 enabled a larger spectrum of possible *E*_*K*_-*V* couples (**Fig. 3B**). Nevertheless, in this system *E*_*K*_ was always larger than *E*_*H*_ (**Fig. 3B**), which would still result in very low cytosolic K^+^ concentrations under otherwise physiological conditions.

**Figure 3.**
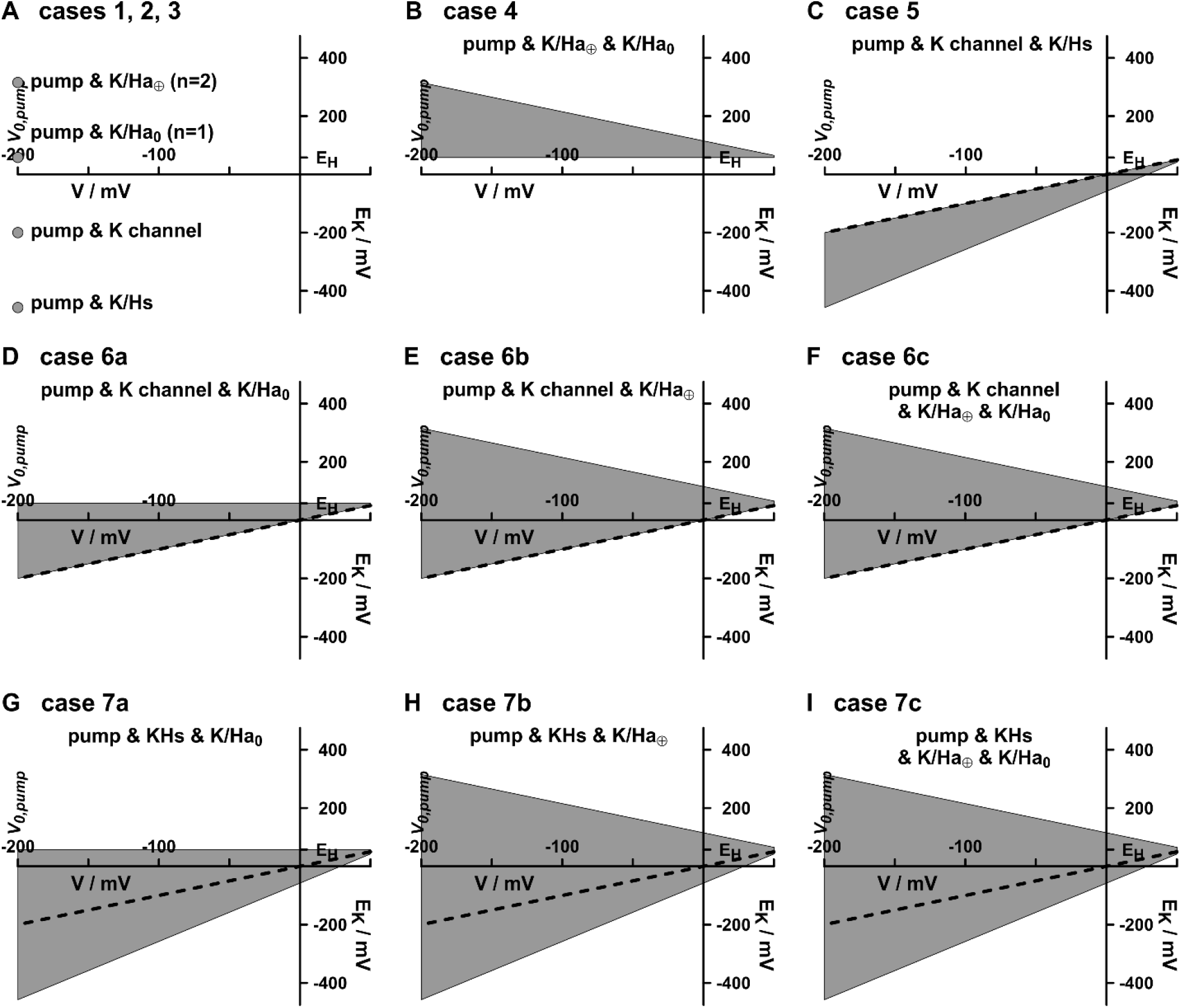
Accessible *E*_*K*_-*V*-ranges for the different K^+^ transporter combinations exemplarily shown for *V*_*0,pump*_ = -200 mV. (A) With a single transporter type only one steady state condition is achievable, which is independent on the transporter activity (grey dots for the different cases). (B-I) The combination of transporter types enables steady states in a wider range (grey areas) by adjusting the activities of the transporters. Nevertheless, there are still limitations. For better orientation, the dashed lines indicate where *E*_*K*_ = *V*. The upper limits in (B, E, F, H, I) are determined by *E*_*K*_ ≤ *n*·*E*_*H*_-(*n*-1)·*V*, in (C) by *E*_*K*_ ≤ *V*, and in (D, G) by *E*_*K*_ ≤ *E*_*H*_. The lower limits in (B) are *E*_*K*_ ≥ *E*_*H*_, in (C, G, H, I) *E*_*K*_ ≥ 2·*V*-*E*_*H*_, and in (D, E, F) *E*_*K*_ ≥ *V*.

### Case 5: K^+^ channels and K^+^/H^+^ symporters

A fifth *‘what-if’* scenario considered the case, in which the proton H^+^-ATPase acted together with K^+^ channels and K^+^/H^+^ symporters. In this system a steady state was established in which the H^+^-ATPase pumped a proton out of the cell, the K^+^/H^+^ symporter allowed the combined influx of a proton and a K^+^ ion, while the channel released a K^+^ ion (**Fig. 2D**). Also here, the combination of different K^+^ transporter types established H^+^ and K^+^ cycles that are energized by the H^+^-ATPase. This drawback, however, was compensated by the flexibility to adjust *V* and *E*_*K*_ by modifying the transporter activity (**Fig. 2E**,**F**). Nevertheless, this flexibility still had its limits. The system considered in case 5 could only achieve a steady state, in which *E*_*K*_ < *V* (**Fig. 3C**).

Under this condition an open potassium channel released K^+^ from the cell, regardless of its gating properties. If in the extreme case the open probability of the K^+^ channel was zero, *i*.*e. p*_*o*_(*V*) = 0, the case 5 converted into the case 2, in which *V* = *V*_*0,pump*_ and *E*_*K*_ = 2·*V*_*0,pump*_–*E*_*H*_.

### Case 6: K^+^ channels and K^+^/H^+^ antiporters

A sixth *‘what-if’* scenario considered the case, in which the proton H^+^-ATPase acted together with K^+^ channels and K^+^/H^+^ antiporters. An electroneutral K^+^/*n*H^+^ antiport was represented by *n* = 1 and an electrogenic antiport by *n* > 1. The system established a steady state, in which the H^+^-ATPase pumped *n* protons out of the cell, while the K^+^/H^+^ antiporter exchanged *n* inflowing protons for an outflowing K^+^ ion, which was reabsorbed by the potassium channel (**Fig. 2G**). *E*_*K*_ and *V* could be flexibly fine-tuned by adjusting the transporter/channel activities (**Fig. 2H**,**I**). Greater antiporter activity increased both *E*_*K*_ and *V*, while greater channel activity increased *V* and decreased *E*_*K*_. However, *E*_*K*_ was always positive of *V* in steady state (**Fig. 3D-F**). With increasing channel activity, the gap between *V* and *E*_*K*_ got smaller, while it got larger with increasing K^+^/H^+^ antiporter activity. If the K^+^/H^+^ antiporter was electroneutral (case 6a) *E*_*K*_ was never larger than *E*_*H*_ (**Fig. 3D**). If it was electrogenic (case 6b), the upper limit of *E*_*K*_ was *n*·*E*_*H*_-*V* (**Fig. 3E**), which is physiologically irrelevant at the plasma membrane. In the physiological range, the main difference between both options was that the maintenance of the H^+^ and K^+^ cycles costed more energy with electrogenic antiporters. The combined presence of electrogenic and electroneutral antiporters (case 6c) did not produce evident new features (**Fig. 3F**) compared to case 6b.

### Case 7: K^+^/H^+^ symporters and K^+^/H^+^ antiporters

A seventh *‘what-if’* scenario considered the case, in which the proton H^+^-ATPase acted together with K^+^/H^+^ symporters and K^+^/H^+^ antiporters of the same type. An electroneutral K^+^/*n*H^+^ antiport was represented by *n* = 1 and an electrogenic antiport by *n* > 1. The system established a steady state, in which the H^+^-ATPase pumped (*n*+1) protons out of the cell, while the K^+^/H^+^ antiporter exchanged *n* inflowing protons for one outflowing K^+^ ion, which was reabsorbed together with a proton by the K^+^/H^+^ symporter (**Fig. 2J**). The behavior of the system in response to changes of the transporter activities was very similar to case 6. *E*_*K*_ and *V* could be flexibly fine-tuned by adjusting the transporter activities (**Fig. 2K**,**L**). Greater antiporter activity increased both *E*_*K*_ and *V*, while greater symporter activity increased *V* and decreased *E*_*K*_. Compared to the cases 5 and 6, however, the system of case 7 gained another level of flexibility. While in case 5 *E*_*K*_ was always smaller than the membrane voltage *V* (**Fig. 3C**) and in case 6 *E*_*K*_ was always larger than *V* (**Fig. 3D-F**), in case 7 *E*_*K*_ could be adjusted in both ranges (**Fig. 3G-I**).

### Case 8: K^+^ channels, K^+^/H^+^ symporters and K^+^/H^+^ antiporters

In an eighth *‘what-if’* scenario, all different K^+^ transporter types were combined. This system showed the total composite complexity of all other systems previously considered, but did not go beyond. *E*_*K*_ and *V* could be flexibly adjusted within the same limits as in case 7 by fine-tuning the transporter activities (**Fig. 4**). This flexibility was again at the cost of energy-consuming K^+^ and H^+^ cycles. For systems with electrogenic K^+^/H^+^ antiporters the energy costs were higher than for systems with electroneutral antiporters.

**Figure 4.**
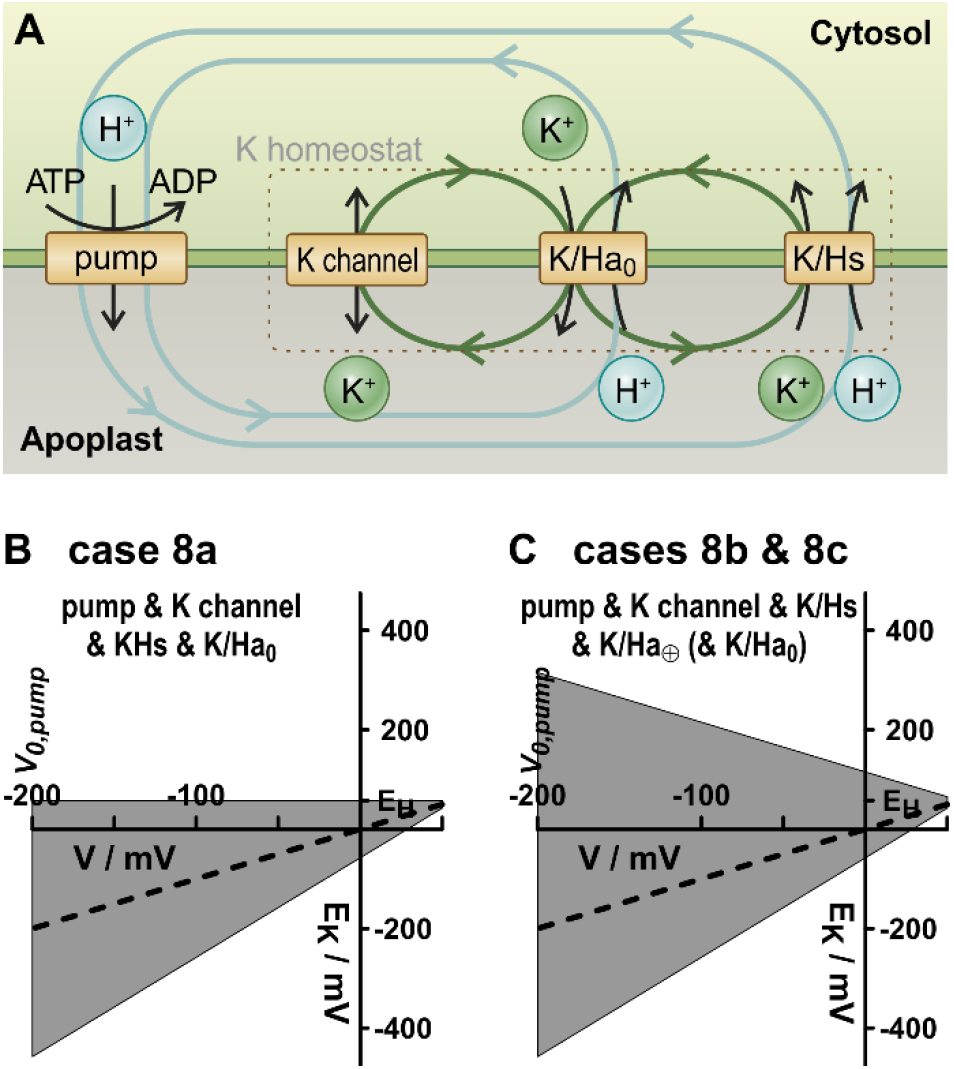
Flexibility of a system with different types of K^+^ transporters. (A) Energized by the H^+^-ATPase, the K homeostat, consisting of K^+^ channels, K^+^/H^+^ symporters and K^+^/H^+^ antiporters, exhibits four cycles that aggregate in two patterns. A proton that is released by the pump is reabsorbed by the antiporter in exchange with a released K^+^, which in turn is reabsorbed (i) either by the channel or (ii) together with another proton by the symporter. (B,C) By adjusting the transporter activity a large range of *E*_*K*_-*V*-steady states can be achieved (grey areas). (B) With electroneutral antiporters the range is limited to *E*_*K*_ ≤ *E*_*H*_. (C) With electrogenic antiporters these limits are further extended to more positive values [*E*_*K*_ ≤ *n*·*E*_*H*_-(*n*-1)·*V*].

### Case 9: Anion channels

After the inventory of the effects of K^+^ transporters, the consequences of charge-balancing anions were considered, first alone and later in combination with the K^+^ transporters. The nineth *‘what-if’* scenario considered the case, in which the proton H^+^-ATPase acted together with anion channels. In such a system, the positive proton pump current was compensated by an efflux of anions (**Fig. S1 A**,**B**). The release of A^-^ led to a negative shift of *E*_*A*_, which also negatively shifted the voltage, at which the H^+^ efflux compensated the A^-^ efflux. A stable steady state could only be reached in this system if *I*_*P*_(*V*) = 0 and *E*_*A*_ = *V*. This means that the membrane voltage adjusted to the steady state voltage of the H^+^-ATPase and that also *E*_*A*_ was fixed to this value causing that there was neither a net A^-^ flux via the channel nor a net H^+^ flux via the pump. Under the conditions presented above (pH_cyt_ = 7.0, pH_apo_ = 6.0; *V*_*0,pump*_ = -200 mV) a steady state condition would be [A^-^]_apo_ = 1 mM / [A^-^]_cyt_ = 3.4× 10^−4^ mM. If other transport processes consumed part of the energy provided by the pump (simulated by setting *V*_*0,pump*_ = -125 mV) the steady state would be [A^-^]_apo_ = 1 mM / [A^-^]_cyt_ = 6.7× 10^−3^ mM, thus still unphysiologically low. As in the cases 1-3 of single K^+^ transporters, the steady state values in case 9 could not be influenced by changing the anion channel activity.

### Case 10: H^+^/A^-^ symporters

A tenth *‘what-if’* scenario considered the case, in which the proton H^+^-ATPase acted together with H^+^/A^-^ symporters. Here, the pumped H^+^ was electrically compensated by a combined 2H^+^/1A^-^ influx (**Fig. S1C**,**D**). While the net proton fluxes were balanced by metabolic/biochemical processes (Sanders and Slayman, 1982; Wegner and Shabala, 2020; Wegner et al., 2021), the uptake of anions shifted *E*_*A*_ positively along the voltage axis and decreased the zero-current voltage of the symporter, *E*_*H/A*_. As a consequence, the membrane voltage, at which the positive and negative electrical fluxes compensated each other, also shifted to more negative values. The system had only one possible steady state at *V* = *V*_*0,pump*_, which implied *E*_*A*_ = 2·*E*_*H*_-*V*_*0,pump*_. Under this condition, the net H^+^/A^-^ flux via the symporter was zero. Considering again the exemplary conditions used above (pH_cyt_ = 7.0, pH_apo_ = 6.0, and *V*_*0,pump*_ = - 200 mV), this system would have equilibrated at [A^-^]_cyt_ = 3× 10^5^ mM for [A^-^]_apo_ = 1 mM. When setting *V*_*0,pump*_ = -125 mV to simulate energy consumption by other transport processes, the gradient would still be [A^-^]_cyt_ = 1.5× 10^4^ mM for [A^-^]_apo_ = 1 mM. Also here, a control of *V* or *E*_*A*_ by varying the H^+^/A^-^ symporter activity was not possible. Thus, symporters alone would generate very high [A^-^]_cyt_ (very positive *E*_*A*_), while anion channels alone would cause very low [A^-^]_cyt_ (very negative *E*_*A*_).

### Case 11: A^-^ channels and H^+^/A^-^ symporters

Next, the combination of anion channels and 2H^+^/A^-^ symporters was tested in the 11^th^ *‘what-if’* scenario (**Fig. 5**). As in the cases of the combinations of potassium transporters, also the presence of different anion transporters in the network allowed the cell now to flexibly adjust *E*_*A*_ and *V* in certain ranges by changing the transporter activities. This gain in flexibility was again at the cost of anion and H^+^ cycles. An ion that was released from the cell was reaccumulated together with two protons via the symporter. To energize this cycle the two protons were pumped out of the cell by the H^+^-ATPase.

**Figure 5.**
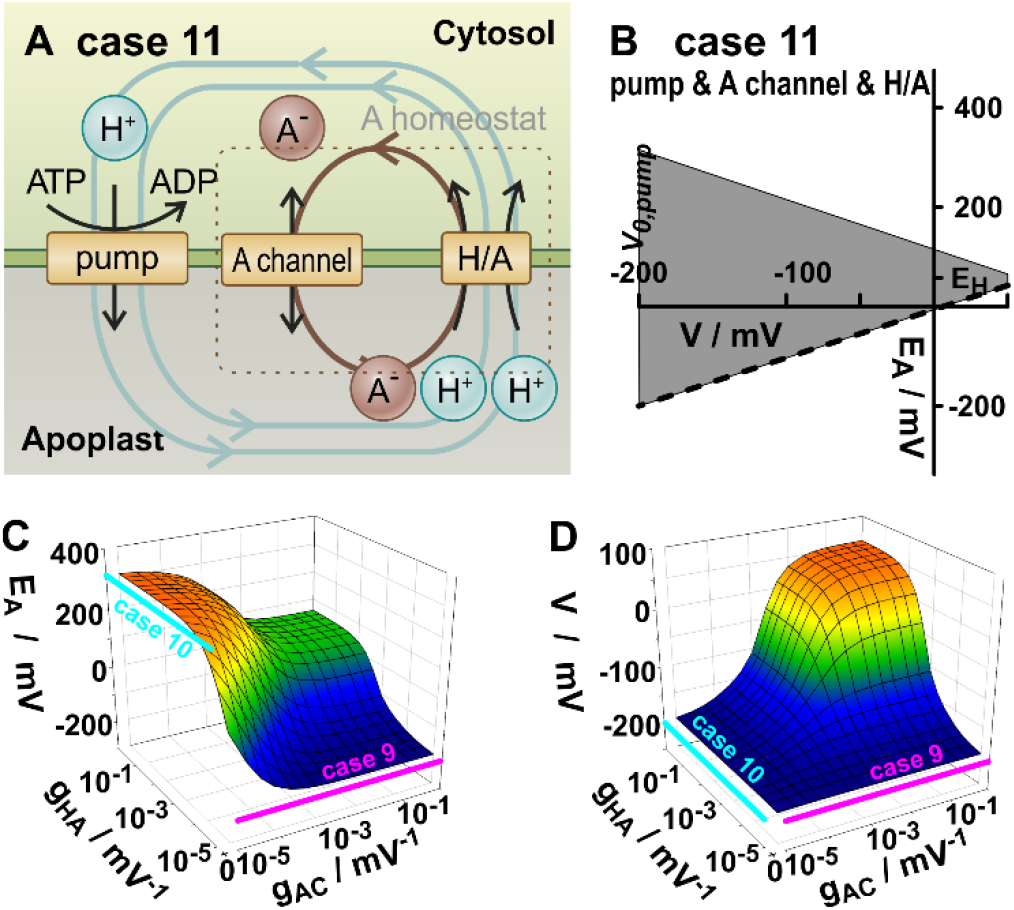
Flexibility of a system with different types of A^-^ transporters (case 11). (A) Energized by the H^+^-ATPase, the A homeostat, consisting of anion channels and electrogenic H^+^/anion symporters, exhibits three cycles. Two protons that are released by the pump are reabsorbed by the symporter together with an anion, which in turn is released by the channel. (B) By adjusting the transporter activity a large range of *E*_*A*_-*V*-steady states can be achieved (grey areas). (C,D) By modifying the transporter activities, the steady state of *E*_*A*_ (C) and *V* (D) can be freely adjusted in a certain range. For comparison, the respective steady states for the systems with only one A^-^ transporter-type: only A^-^ channels (*g*_*HA*_ = 0; Case 9, magenta lines) and only 2H^+^/1A^-^ symporters (*g*_*AC*_ = 0; Case 10, cyan lines).

### Cases 12 to 16: One homeostat in combination with a single transporter

To understand how the combination of K^+^ and A^-^ transporters effected the homeostatic control of the respective other ion, the two anion transporters (“A-homeostat”) were combined with the pump and with one of each of the three K^+^ transporter types (case 12: K^+^ channel, case 13: K^+^/H^+^ symporter, case 14: K^+^/H^+^ antiporter). The “K-homeostat” (K^+^ channel, K^+^/H^+^ symporter and 1K^+^/1H^+^ antiporter) was combined with the pump and the anion channel (case 15), or the H/A symporter (case 16; **Fig. S2**). In all these cases, the respective homeostat determined *V*, the steady state membrane voltage and the particular concentration gradient (*E*_*K*_ for K-homeostat, *E*_*A*_ for A-homeostat). Varying the activity of the single transporters had neither influence on those, nor had this any influence on the concentration gradient of the ion they transport (**Fig. S2**). The *E*_*K*_ and *E*_*A*_ values, respectively, were fixed in steady state to the prevailing membrane voltage as shown in cases 1-3, 9, and 10. In the cases 12-16, the cycles of the respective homeostat consumed part of the electrochemical energy provided by the H^+^-ATPase, which manifested in the difference between *V* and *V*_*0,pump*_. It may be remembered that in some examples for the cases 1, 2, 3, 9, and 10 the same effect was simulated by modifying *V*_*0,pump*_ to clamp a different membrane voltage.

### Case 17: K-homeostat and A-homeostat

When instead of single transporters, both homeostats were combined (case 17, **Fig. S3**) the system gained the full flexible range. Variations in the activity of a K^+^ transporter affected not only the steady state of *E*_*K*_ but also the membrane voltage *V* and, via *V*, the steady state of *E*_*A*_. In the same way, variations in the activity of an anion transporter directly affected both *E*_*A*_ and *V* and, via *V*, also *E*_*K*_. Nevertheless, the orientation of *E*_*K*_ towards *V* was exclusively determined by the K transporters while the orientation of *E*_*A*_ towards *V* was only fixed by the anion transporters. In other words, only the corresponding homeostat was able to set whether (i) *V*-*E*_*X*_ > 0, (ii) *V*-*E*_*X*_ = 0, or (iii) *V*-*E*_*X*_ < 0. Variations in the other homeostat could not cause a switch between these three possibilities.

### Cases 18-20: K-homeostat and sugar transporters

After having systematically analyzed transporter networks for cations and for anions alone and in combination, as next the transport of an electrically neutral nutrient was elaborated. For this purpose, the H^+^-ATPase and the K-homeostat (comprising K^+^ channels, K^+^/H^+^ symporters and electroneutral K^+^/H^+^ antiporters) were combined in a 18^th^ *‘what-if’* scenario with proton-coupled sugar transporters (HC), in a 19^th^ scenario with sugar uniporters (SWEETs) and in an 20^th^ with both sugar transporter types (**Fig. 6**). For simplicity, the A-homeostat was not included. As previously shown, its presence would influence the transport of K^+^ and sugar only by consuming part of the cellular energy, which manifests itself in a different steady state value of *V*. Such an effect could also be simulated by other processes, such as adjusting *V*_*0,pump*_ or allowing the K-homeostat to consume more energy.

**Figure 6.**
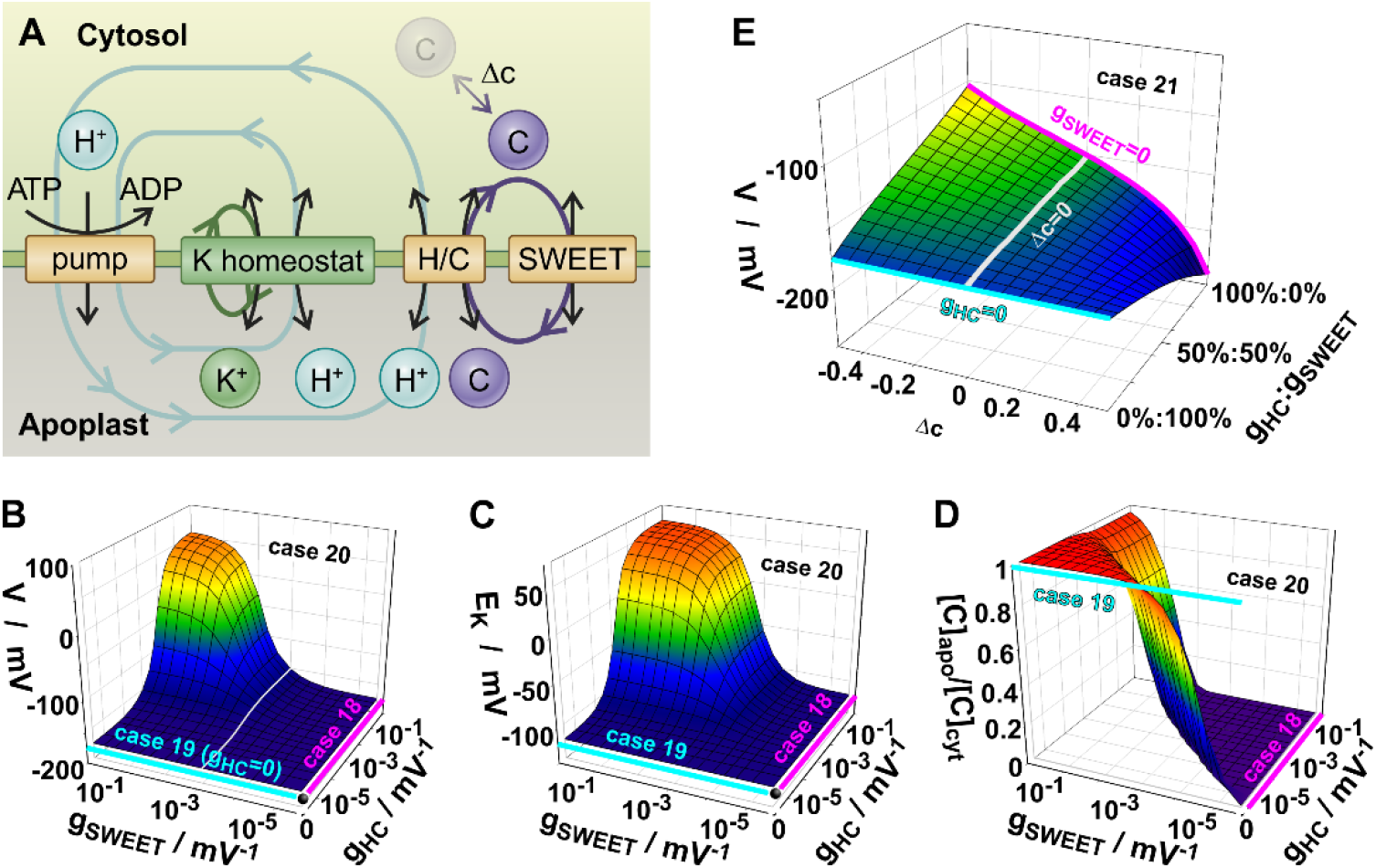
Combination of K^+^ and sugar transport. The K-homeostat (Fig. 4) was combined with sugar transport via uniporters (SWEET) and/or proton-coupled symporters. (A) In addition to the cycles of the K-homeostat the presence of both sugar transporter types established proton/sugar cycles, in which a sugar molecule is released by SWEET and reabsorbed by the symporter together with a proton that is released by the pump. Additionally, the metabolization of sugar was also simulated (Δc). (B-D) When both sugar transporters are present (case 20), the membrane voltage (B) and the transmembrane sugar gradient (D) can be adjusted by fine-tuning the transporter activities. The altered energetic status of the cell indicated by the steady state voltage *V* (B) also affected *E*_*K*_ (C). When only one sugar transporter type is present (case 18: only H/C; case 19: only SWEET) the activity of the transporter does not affect *V, E*_*K*_ or the sugar gradient in steady state (magenta and cyan lines). (E) Case 21. Effect of metabolic sugar production (Δc > 0) or sugar consumption (Δc < 0) on the steady state membrane voltage *V*. An effect was only observable in the presence of electrogenic H/C transporters. If only electroneutral SWEETs were present, sugar metabolism did not affect *V* (cyan line).

To illustrate the effects of the sugar transporters, the different parameters of the K-homeostat were set to achieve a steady state of *V* ≈ -175 mV and *E*_*K*_ ≈ -117 mV in the absence of any sugar transporter and were kept constant in all scenarios 18-20. Different parameters would not have changed the results qualitatively. In the presence of proton-coupled sugar transporters alone (case 18), *V* and *E*_*K*_ did not change irrespective of the activity of the H/C transporter (**Fig. 6A-D**, case 18, magenta). Also the sugar concentration gradient was independent of whether H/C was moderately or highly active. The steady state value was always [*C*]_*cyt*_ ≈ 1.1×10^4^·[*C*]_apo_, and was determined by the pH gradient and the membrane voltage [ln([*C*]_*apo*_/[*C*]_*cyt*_)=*F*/(*RT*)·(*V*-*E*_*H*_)]. When the proton-coupled sugar transporter was replaced by a sugar uniporter (case 19), this steady state gradient collapsed to [*C*]_*cyt*_ = [*C*]_apo_. Also in this case, neither the sugar gradient, nor *E*_*K*_, nor *V* were affected by changing the uniporter activity (**Fig. 6A-D**, case 19, cyan). If, however, both transporter types were simultaneously in the membrane, the system became sensitive towards changes in the transporter activities. The presence of both transporter types established an apparently futile C-cycle that consumed cellular energy. As a consequence, *V* increased with increasing sugar cycling and this reduced driving-force for ions affected *E*_*K*_ as well (**Fig. 6A-D**, case 20). Thus, the adjustment of the sugar gradient by fine-tuning the activity of the sugar transporters coupled back to K^+^ homeostasis via the energy status of the cell. Interestingly, the same thought experiments with only one active K^+^ transporter at a time reproduced the cases 1 (*E*_*K*_ = *V*), 2 (*E*_*K*_ = 2·*V* – *E*_*H*_), and 3 (*E*_*K*_ = *E*_*H*_), respectively. There, the steady state voltage *V* was less negative the more energy the sugar cycle consumed. Therefore, to adjust K^+^ and sugar gradients independent of each other, for each nutrient two different transporter types were needed in the membrane, resulting consequently in nutrient cycles for K^+^ and sugar.

### Case 21: Metabolism can affect the energetics of membrane transport

In the same system as before, the influence of metabolic processes on the homeostatic conditions was tested. For this purpose, the production (Δc > 0) or consumption (Δc < 0) of cytosolic sugars was simulated (**Fig. 6A**). Without any sugar transporter in the membrane, the K-homeostat was not affected at all by metabolic changes of the sugar concentration. Similarly, when only electroneutral sugar uniporters were present, sugar production or consumption affected only the transmembrane sugar gradient but had no effect on the steady state membrane voltage *V* (**Fig. 6E**, cyan) or *E*_*K*_. However, in the presence of electrogenic proton-coupled sugar transporters, sugar metabolism affected the membrane voltage (**Fig. 6E**) and, via this, *E*_*K*_. Compared to the conditions without metabolism, sugar production created an additional sugar gradient out of the cell, which resulted in an additional efflux via sugar uniporters and sugar-proton symporters. While the electroneutral transport via uniporters did not affect the membrane voltage, the electrogenic transport via proton-coupled transporters provoked a more negative *V*. Conversely, sugar consumption led to an additional sugar gradient into the cell, which resulted in an additional sugar influx. When mediated by proton-coupled symporters, this influx provoked a less negative *V* (**Fig. 6E**). Thus, if metabolic processes are coupled to electrogenic transport processes, they can interfere with general homeostatic control.

### Generalization for more nutrients

In further iterative steps the transporter system could be brought successively closer to the real physiological situation. K^+^ transporters and sugar transporters could be combined into a new “K/C homeostat” which was then combined with transporters for another nutrient or ion. The conclusions drawn before for sugar transport were then valid for the new substrate. The different homeostat modules were largely independent of each other but coupled via the energetic status of the cell. A special case that needs to be investigated in a separate study, however, would be flux coupling in one transporter as may be the case with Na^+^ and K^+^ in HKT channels (Riedelsberger et al., 2019; Riedelsberger et al., 2021).

### Cases 22: Symplastic diffusion

All the scenarios so far considered a static isolated cell. The reality in a plant, however, is more complex. For instance, cells are connected to neighbors via plasmodesmata enabling diffusion. This process could be covered in the model by a mathematical description similar to case 21. A loss of nutrients by diffusion to a neighboring cell was equivalent to a transport process out of the cell, while a diffusive gain of nutrients was equivalent to a transport process into the cell. For exemplary illustration, the transporter network of case 12 (**Fig. S2A**) was considered again in a 22^nd^ *‘what-if’* scenario extended with these diffusion processes (**Fig. S4**). Without diffusion *E*_*K*_ was identical to the steady state membrane voltage irrespective of the activity of the K^+^ channel. Likewise, *E*_*A*_ was constant in all these conditions. Diffusion out of the cell did not markedly influenced *V*, caused a moderate decline of *E*_*A*_, but strongly affected *E*_*K*_ provoking *E*_*K*_ > *V*. The latter effect was more pronounced the lower the K^+^ channel activity was; thus, the less the loss by diffusion could be compensated by uptake. Diffusion into the cell had the opposite effects. Diffusion could therefore be considered as an additional transport process with which a cell gained flexibility to adjust *E*_*K*_ even with a single membrane transporter type for K^+^, as a K^+^ channel in this case. In a normal physiological condition, however, *g*_*KC*_ is in the order of magnitude of 10^−1^ mV^-1^ suggesting that the influence of diffusion on the homeostatic conditions is not strong. Additionally, if the diffusion process stops because plasmodesmata close or because a symplastic concentration gradient is missing, even a symplastically connected cell is not different from the cases 1-21 with all the consequences described above.

### Case 23: Volume changes

In principle, also volume changes could affect homeostatic processes. During stomal movement, for instance, a guard cell increases its volume by ∼25% and its surface area by ∼15% (Meckel et al., 2007; Jezek and Blatt, 2017). Compared to a static cell as it was considered so far, the increased volume would imply a dilution of the cytosolic nutrient concentrations and the increased surface area would imply an increased electrical capacitance of the membrane, which in turn would result in a reduction of the magnitude of the membrane voltage. To estimate the impact of these effects, they were included in the mathematical description (Supplementary Equations S1 and S2) and were exemplarily simulated in a 23^rd^ *‘what-if’* scenario (**Fig. S5**), which is based on cases 12 and 22. To estimate the parameter range, a guard cell was considered that changed rather rapidly in a linear manner from 4 pl and 20 pF (closed) to 5 pl and 23 pF (open) in 60 min (Hosy et al., 2003). From these values and with exemplarily chosen *V* ≈ - 100 mV, *I*_*max*_ = 20 pA, and [K^+^]_*cyt*_ = 100 mM, the orders of magnitude of γ = -*V*·*I*_*max*_/·*dC*/*dt* ≈ 4×10^−6^ and Δ*k* = -*e*_*0*_/*I*_*max*_·*N*_*A*_·[K^+^]_*cyt*_·*dVol*/*dt* ≈ -0.15 were calculated. Slower stomatal movements would result in even lower values. In comparison to the membrane transport processes (*e*.*g*., a usual value for *i*_*p*_, eqn. S1, is in the range of 0.2 - 0.7) the contribution of the parameter γ is neglectable. The change of the volume, however, could influence the homeostatic conditions (**Fig. S5B**) similar to case 22. Nevertheless, each guard cell will sooner or later reach its maximal or minimal expansion (*dVol*/*dt* = 0) and for these conditions the cases 1-21 also apply to guard cells.

### The role of organelles in homeostasis

Up to now, the problem of homeostasis has been addressed at a single membrane. The presented results were exemplarily obtained for the plasma membrane. But they can be adapted in a similar manner to the vacuolar membrane or any other organelle membrane (**Fig. 7A**). In the cellular context, however, these membranes are not isolated. Instead they communicate with each other via the cytosolic concentrations (Vincent et al., 2017; Horaruang et al., 2020). Previous studies have shown that membrane sandwiches exhibit another level of dynamics (Gajdanowicz et al., 2011; Schott et al., 2016; Dreyer et al., 2017; Dreyer et al., 2019). Therefore, nutrient homeostasis was also investigated with respect to these dynamics. It turned out that cytosolic concentrations could remain equilibrated in two different modes. In a long-term stable steady state, the net-fluxes of each substrate were zero across each membrane separately (**Fig. 7A**), while on a much shorter time-scale the shuttling of nutrients from the lumen of an organelle to the apoplast and/or vice versa was possible as well (**Fig. 7B-E**). The first mode implied the presence of energy consuming nutrient cycles. The second mode may function without such cycles. However, it was not stable in the long-term because the nutrient net-fluxes changed the nutrient concentrations in particular in the lumen which is an enclosed compartment without additional inlets and outlets. Also the concentrations in the apoplast could change. However there, diffusion processes may partially mitigate this effect. Importantly, the changed concentrations reduced the driving forces for the fluxes. In the end, every possible arrangement of the second mode (**Fig. 7B-E**) resulted in the long-term stable steady state of the first mode (**Fig. 7A**). Thus, the processes investigated in this study are not special cases, but a fundamental basis of nutrient homeostasis in plant cells.

**Figure 7.**
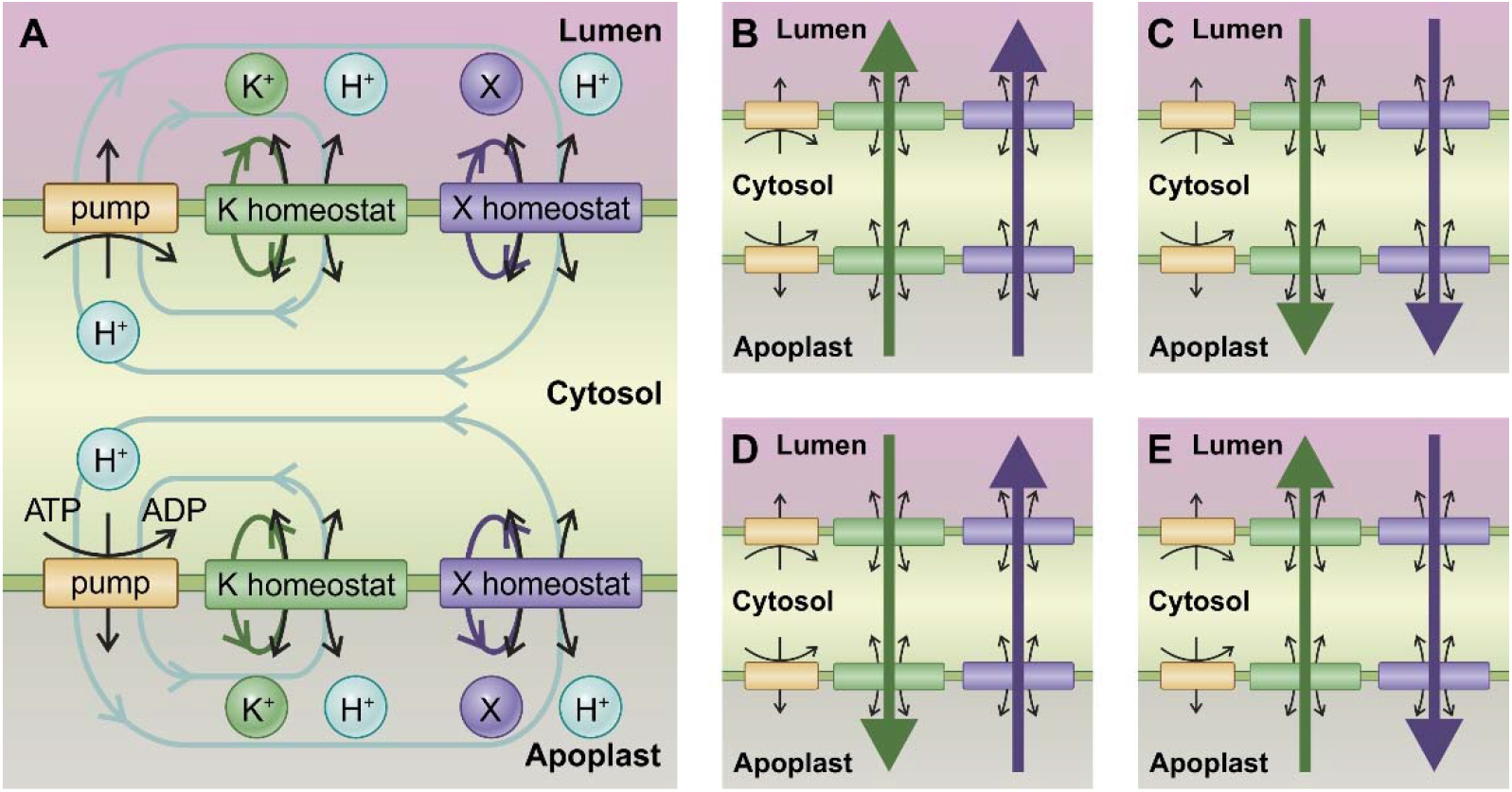
Combination of homeostats in membrane sandwiches. (A) Homeostats at the plasma membrane and at organelle membranes may maintain the cytosolic nutrient concentrations constant for longer time by independently cycling the nutrients across the different membranes. By adjusting the transporter activities, the concentrations and membrane voltages can be fine-tuned. (B-E) Nutrients can be sequestrated into or remobilized from organelles by shuttling them across the cytosol. For two different nutrients the four possibilities of nutrient fluxes are shown. These systems are only temporarily in flux-steady state as the fluxes change the concentrations in the lumen and the apoplast while cytosolic concentrations remain stable. On longer time-scales the systems (B-E) approach a situation displayed in (A).

## Discussion

Transport processes across membranes are ruled by fundamental, well-understood laws. This allows to describe them in mathematic terms (Beilby, 2007; Hills et al., 2012; Beilby and Al Khazaaly, 2016; Dreyer, 2017) and to run computational cell biology simulations. Such solidly based thought experiments have an inestimable value in gaining new insights because they enable to test conditions that are hard to achieve in conventional wet-laboratory experiments. The present study analyzed the basis of homeostasis and allowed far-reaching conclusions for our physiological understanding.

### Two different transporter types are needed for homeostatic control

Comparison of cases 1-3 with 5-8, of cases 9 and 10 with 11, and of cases 18 and 19 with 20 showed that a cell gained flexibility by combining at least two different transporter types. Only then the different *E*_*X*_ and *V* could be adjusted in certain ranges by fine-tuning the transporter activities. This insight may explain why plants have different transporter types for many macro nutrients. It is not that these transporters have different ‘affinities’ and act in different concentration ranges (Dreyer and Michard, 2020). Instead, they act together and simultaneously, and thus can control the transmembrane concentration gradient, in order to keep the cytosolic concentration constant even under varying external conditions. An important feature in this mechanism is that the transport through these transporter types is energized differently. For instance, one type transports along the electrochemical gradient of the nutrient while others use coupled nutrient and proton gradients as driving force. Only the combination of these differently energized transport processes enables flexible control. In this context, also other processes like diffusion or metabolism may be considered as “transporter-equivalent”. As indicated in scenarios 21, 22 and 23 such a process can replace under certain conditions one of the two necessary transporter types.

### ’Futile’ nutrient cycles in plants are important for homeostasis

The flexibility to adjust *V* and *E*_*X*_ by two transporter types was inevitably linked to the presence of apparently futile nutrient cycles. Thus, these ‘futile’ cycles are not as futile as they seem. Instead, they are important for the dynamic flexibility of the system. Nutrient cycling is a widely observed phenomenon in plants (Britto and Kronzucker, 2006), not only in normal conditions but in particular also under salt stress (Munns et al., 2020; Shabala et al., 2020). Such cycles are often considered as an avoidable energy dissipation that could be eliminated in breeding programs. The new fundamental insights provided here may set a new spotlight on this issue. The elimination of ‘futile’ cycles by silencing transporters will straightforwardly reduce the ability of the plant for homeostatic control and will most likely come at the expense of fitness. The proton pump together with the metabolically determined transmembrane pH gradient provide strong steady driving forces for transport processes in plant cells. The examples presented in this study indicate that for a plant cell a severe problem is not necessarily the uptake of a nutrient but its over accumulation, in particular if it is a macronutrient. Nutrient cycles are a very elegant way out of this dilemma. They do not only allow the tight control of the cytosolic nutrient concentration but also allow to dissipate excess energy if needed without having to shut down cellular energy production at greater expense.

### Alternative scenarios

The presented cycling scenario for K^+^ homeostasis is a simple, rather intuitive conclusion from experimentally determined properties of membrane transporters. It should be mentioned, however, that in particular for guard cells, an average constant cytosolic [K^+^] has been explained with a more complex alternative scenario. It was proposed that guard cells are unlikely to be found in a state in which the net flux of an osmotically active solute is zero. Instead, it was suggested that they transit between situations of osmotic solute uptake and loss, to achieve by time-averaged approximation a dynamic range of (quasi-)steady states in solute contents (Thiel et al., 1992; Chen et al., 2012). The conclusions were drawn from the observations on excised guard cells in epidermal strips that the membrane voltage can spontaneously oscillate in an action potential-like behavior with periods from a few tens of seconds to many minutes (Thiel et al., 1992; Gradmann et al., 1993) between two quasi-stable states, a depolarized state associated with K^+^ and Cl^-^ efflux, and a hyperpolarized state at which ion uptake is supposed to occur. Intriguingly, these spontaneous oscillations were not observed in guard cells in intact plants although these cells can stay over longer time periods in the hyperpolarized state (Roelfsema et al., 2001; M.R.G. Roelfsema, personal communication) rising the questions of how guard cells achieve K^+^ homeostasis *in vivo*, in particular when they are maximally swollen, without oscillations. K^+^ cycling mediated by the K-homeostat (case 8) may provide an answer to this question.

### K^+^ homeostasis and electrical signaling

Instead of being a mechanism of homeostasis, the observed membrane potential oscillations in excised guard cells may be related to other physiological processes like action potentials (APs) which are essential parts of electrical signaling in plants (Trebacz et al., 2006; Choi et al., 2016; Hedrich et al., 2016; Cuin et al., 2018; Gilroy et al., 2018; Farmer et al., 2020). Essential for APs is that excitable cells last over a long period at a hyperpolarized resting voltage (Böhm et al., 2016; Iosip et al., 2020). Each spontaneous oscillation to release over-accumulated osmotically active ions, as K^+^ and Cl^-^, would fire an action potential and would strongly distort electrical signaling. In the Venus flytrap (*Dionaea muscipula*), for instance, such spontaneous oscillations would cause false triggering of the trap mechanism (Scherzer et al., 2019). A trigger hair cell in Dionaea, as an example for an excitable cell, has to rest in a steady-state with a reasonable negative membrane voltage until it gets mechanically stimulated to fire an action potential. The resting membrane voltage *V* is determined by the activity of all ion transporters in the membrane and is in the range of -200 mV < *V* < -100 mV (Scherzer et al., 2019; Iosip et al., 2020). Considering that K^+^ and anions like Cl^-^ are the main players in the execution of plant APs (Beilby and Coster, 1979; Beilby and Walker, 1996; Beilby and Al Khazaaly, 2016; Beilby and Al Khazaaly, 2017; Cuin et al., 2018; Kisnieriene et al., 2019) such a condition could be achieved by a transporter network of case 17 or 12. While case 17 can flexibly realize all three possibilities (i) *V* < *E*_*K*_, (ii) *V* = *E*_*K*_, or (iii) *E*_*K*_ < *V*, case 12 is limited to *V* = *E*_*K*_. In specific circumstances, with stable external [K^+^] conditions, this could be sufficient. However, the limited flexibility would certainly be the weak point of such a system.

### Conclusion

The comprehensive analysis of the dynamics of transporter systems suggests that nutrient cycles are important for homeostatic control. The elimination of these cycles by silencing transporters will likely decrease the fitness of the plant. To improve nutrient use efficiency in crops, different strategies are needed. One path could be the continuation with computational cell biology simulations and dry-lab experiments. This might provide further fundamental insight into the

dynamics of the systems and may preserve us from too many dead-end strategies to be tested in the wet-lab and in field trials.

## Materials and Methods

### Mathematical Description of Transporter Activities

#### H^+^-ATPase

The voltage-dependent current of the H^+^-ATPase was described with a mechanistic 6-state pump model (Dreyer, 2017; Reyer et al., 2020) that pumps with a 1 H^+^ : 1 ATP ratio (Blatt et al., 1990):

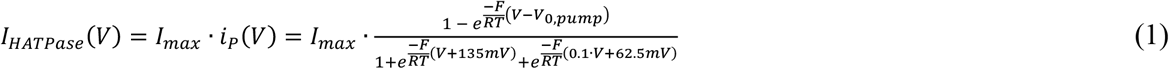

Here *F* is the Faraday constant, *R* the gas constant and *T* the absolute temperature. *I*_*max*_ is the maximum current, which depends on the number of active pumps in the membrane and the cytosolic and apoplastic proton concentrations. *V*_*0,pump*_ is the voltage, at which the pump current is zero. It depends on the cytosolic and apoplastic proton concentrations and on the cytosolic ATP, ADP, and Pi concentrations, *i*.*e*., on the energy status of the cell (Rienmüller et al., 2012). In the examples presented in this study, this value was set exemplarily to *V*_*0,pump*_ = -200 mV; different values would not change the results qualitatively. In some examples, this parameter was modified to simulate the consumption of energy by other transport processes.

#### 1 H^+^ : 1 K^+^ symporter

Proton (*J*_*H,KHs*_) and potassium net flux (*J*_*K,KHs*_) from the cell and the current-voltage relationship

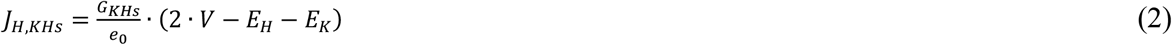

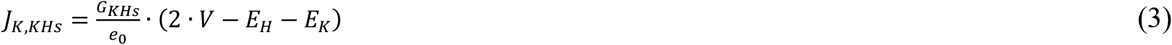

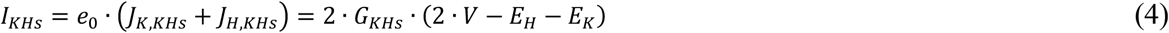

where *V* is the membrane voltage, *e*_*0*_ the elementary charge, *E*_*K*_ the Nernst voltage for potassium and *E*_*H*_ the Nernst voltage for protons. *G*_*KHs*_ is the membrane conductance of this transporter type (unit pA/mV).

#### n H^+^ : 1 K^+^ antiporter

Proton (*J*_*H,KHa*_) and potassium (*J*_*K,KHa*_) net efflux and the current-voltage relationship (*I*_*KHa*_) of an antiporter that transports *n* protons for one K^+^ ion:

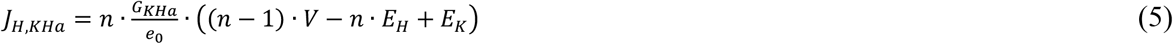

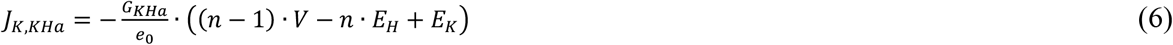

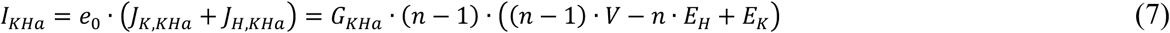

with *G*_*KHa*_ being the membrane conductance of this transporter type. In analogy to all other transporters, also for the electroneutral 1:1 antiporter (*n* = 1) a parameter *G*_*KHa0*_ with the unit of an electrical conductance (pA/mV) can be defined, although this transporter does not conduct electrical current. This trick has technical reasons to unify the equations. It is evident for *n* = 1 that *I*_*KHa0*_ = 0 and that the proportional factor for the fluxes *G*_*KHA0*_/*e*_*0*_ has the unit mV^-1^.

#### Voltage-gated K^+^ channel

Potassium net efflux (*J*_*K_KC*_) and the current-voltage relationship (*I*_*KC*_) of a voltage-gated potassium channel:

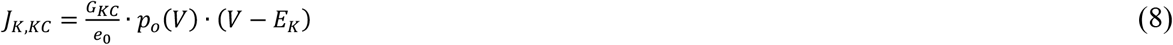

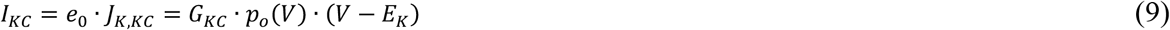

with *G*_*KC*_ (unit pA/mV) being the membrane conductance for this channel type. The factor *p*_*O*_(*V*) describes the voltage-dependent probability that a channel is open. A suitable mathematical description for hyperpolarization-activated K^+^ channels is (Brüggemann et al., 1999):

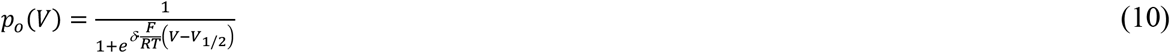

with δ = 1.6 and *V*_*1/2*_ = -175mV. Different values for the gating charge or the half-maximal activation voltage in the physiologically reasonably intervals (δ = 0.5 … 2; *V*_*1/2*_ = -100 mV … -200 mV) did not change qualitatively the results presented in this study.

#### Anion channel

Anion net efflux (*J*_*A_AC*_) and the current-voltage relationship (*I*_*AC*_) of an anion channel of the SLAC1/SLAH-type:

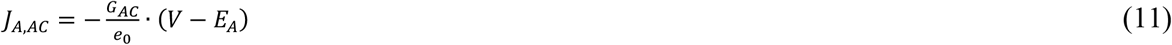

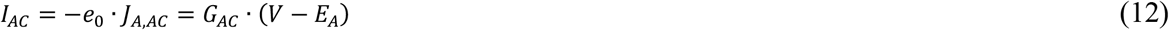

with *E*_*A*_ being the Nernst voltage of the anion and *G*_*AC*_ (unit pA/mV) the membrane conductance for this channel type.

#### 2 H^+^ : 1 A^-^ symporter

Proton (*J*_*H,HA*_) and anion (*J*_*A,HA*_) net efflux and the current-voltage relationship (*I*_*HA*_) of a 2:1 H^+^-coupled H^+^/A^-^ symporter:

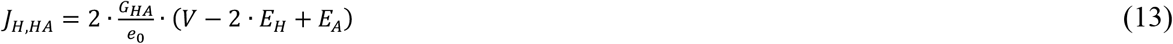

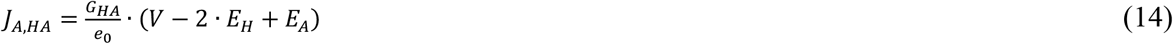

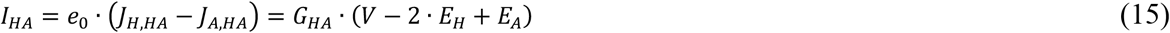

where *G*_*HA*_ is the membrane conductance of this transporter type (unit pA/mV).

#### Sugar uniporter

Sugar net efflux (*J*_*C,SWEET*_) of a sugar uniporter (SWEET type):

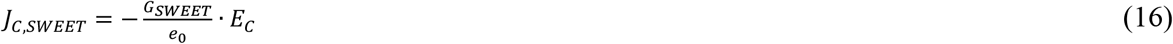

with 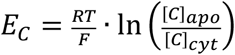 and the apoplastic and cytolic sugar concentrations [C]_apo_ and [C]_cyt_.

Although an electroneutral sugar transporter does not have an electrical conductance, *G*_*SWEET*_ and *E*_*C*_ were defined in analogy to the other transporters to unify the mathematical description.

#### Proton-coupled sugar transporter

Proton (*J*_*H,HC*_) and sugar (*J*_*C,HC*_) net efflux and the current-voltage relationship (*I*_*HC*_) of a 1:1 H^+^/C symporter (Carpaneto et al., 2005):

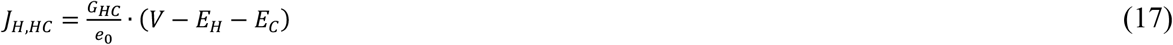

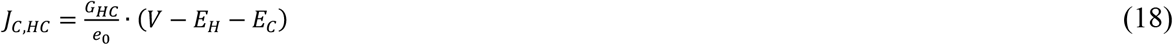

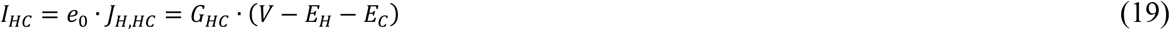

with the membrane conductance of this transporter type, *G*_*HC*_ (unit pA/mV).

### Changes in voltage and concentrations and steady state conditions

The membrane voltage changes according to

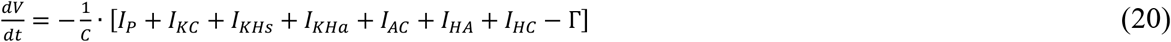

the cytosolic potassium concentration according to

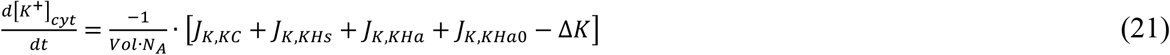

the cytosolic anion concentration according to

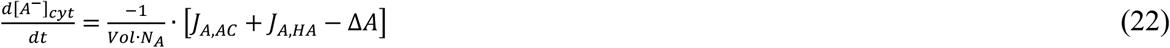

and the cytosolic sugar concentration according to

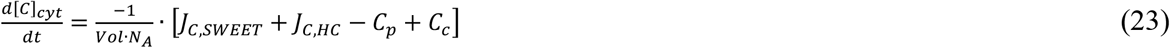

with the membrane capacitance *C*, the cellular volume *Vol*, and the Avogadro constant *N*_*A*_. The parameters Δ*K*, Δ*A, C*_*p*_, and *C*_*c*_ describe concentration changes due to metabolic processes (*e*.*g*., photosynthesis, *C*_*p*_, and respiration, *C*_*c*_, in the case of sugar) or diffusion into (Δ*K* = Δ*A* > 0) and out of (Δ*K* = Δ*A* < 0) the cell, *e*.*g*. via plasmodesmata. In special cases, the parameters Δ*K*, Δ*A*, and ΔC = *C*_*p*_ - *C*_*c*_ may also be considered together with Γ to simulate concentration and voltage changes due to changes in the cellular volume and membrane surface (*e*.*g*., guard cell swelling and shrinking). For more details see Supplementary Equations S1 and S2. The steady state analyzed in this study is defined by the conditions:

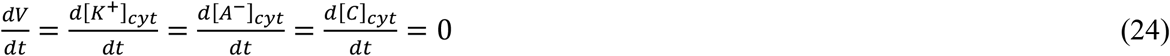

### Proton concentrations

Proton concentrations are predominantly determined by metabolic processes (Sanders and Slayman, 1982; Wegner and Shabala, 2020; Wegner et al., 2021). Therefore, *E*_*H*_ was fixed to +57.6 mV for the steady state analyses in this study. This value represented a pH gradient of pH_cyt_ = 7.0 and pH_apo_ = 6.0. Other physiological pH gradients would not change the presented results qualitatively.

### Renormalization to eliminate a redundant parameter

Equations (20-23) were multiplied by the factor 1 = *I*_*max*_/*I*_*max*_. *I*_*max*_ was combined with the pre-factors to *I*_*max*_/*C* and *I*_*max*_/(*Vol*·*N*_*A*_), while 1/*I*_*max*_ was used to normalize the parameters to the maximal pump current: *i*_*P*_(*V*) = *I*_*P*_(*V*)/*I*_*max*_ (without unit), *g*_*X*_ = *G*_*X*_ /*I*_*max*_ (X=KHs, KHa, KHa0, KC, AC, HA, SWEET, HC; all *g*_*X*_ with unit mV^-1^), *c*_*p*_ = *C*_*p*_·*e*_*0*_/*I*_*max*_, *c*_*c*_ = *C*_*c*_·*e*_*0*_/*I*_*max*_, Δ*c* = *c*_*p*_ - *c*_*c*_, Δ*k* = Δ*K*·*e*_*0*_/*I*_*max*_, Δ*a* = Δ*A*·*e*_*0*_/*I*_*max*_, and γ = Γ/*I*_*max*_ (all without unit). This renormalization eliminated one redundant parameter and yielded the final equation system for the steady state conditions (Supplementary Equation S3).

### Numerical determination of steady state conditions

For each parameter set *g*_*x*_ the equation system has a steady state solution for the membrane voltage *V* and the *E*_*X*_-values. These were determined first by solving eqn (S3) and finally by resolving numerically the resultant implicit equations with self-made scripts.

### Modell assessment

It should be noted that all conclusions drawn in this study are independent of the exact description of the fluxes *J*_*X*_ (**Fig. S6**). The screening of the entire reasonable parameter space [0,∞) for the *g*_*X*_ guaranteed the coverage of all possible cases that a cell can achieve in the considered scenarios and allowed to assess the limits of the flexibility of the system under consideration in steady state even without prior knowledge of specific transporter features.

## Supplemental Material

Equations S1 and S2: Mathematical representation of concentration and voltage changes in swelling or shrinking cells.

Equation S3: Mathematical representation of the steady state conditions of the systems considered in this study.

Figure S1: Cases 9 and 10. Combination of single anion transporters with the H^+^ pump.

Figure S2: Cases 12 to 16. Combination of homeostats with a single transporter for another ion species.

Figure S3: Case 17. Combination of K- and A-homeostats.

Figure S4: Consequences of diffusion through plasmodesmata.

Figure S5: Consequences of changes in cell size.

Figure S6: Equivalence of nonlinear modeling and linear modeling.

## Acknowledgments

The author thanks Rob Roelfsema, Würzburg, for helpful discussions regarding membrane potential oscillations in guard cells.

